# Assessing genotype adaptability and stability in perennial forage breeding trials using random regression models for longitudinal dry matter yield data

**DOI:** 10.1101/2023.03.20.533513

**Authors:** Claudio Carlos Fernandes Filho, Sanzio Carvalho Lima Barrios, Mateus Figueiredo Santos, Jose Airton Rodrigues Nunes, Cacilda Borges do Valle, Liana Jank, Esteban Fernando Rios

## Abstract

Genotype selection for dry matter yield (DMY) in perennial forage species is based on repeated measures over time. Repeated measurements in forage breeding trials generate longitudinal datasets that must be properly analyzed giving a useful interpretation in the genotype selection process. In this study, we have presented a random regression (RRM) approach for selecting genotypes based on longitudinal DMY data generated from ten breeding trials and three perennial species, alfalfa (*Medicago sativa* L.), guineagrass (*Megathyrsus maximus), and* brachiaria (*Urochloa spp*.). We also proposed the estimation of adaptability based on the area under the curve and stability based on the curve coefficient of variation. Our results showed that RRM always approximated the (co)variance structure into an autoregressive pattern. Furthermore, RRM can offer useful information about longitudinal data in forage breeding trials, where the breeder can select genotypes based on their seasonality by interpreting reaction norms. Therefore, we recommend using RRM for longitudinal traits in breeding trials for perennial species.

## Introduction

Genotype selection in perennial forage species involves multiple assessments conducted repeatedly over time, across various seasons and years. Therefore, the evaluation of multi-harvest forage breeding trials is time-consuming and expensive. In this context, proper statistical methods that accurately predict the true genotypic values is crucial [1]. As the genotypes experience different environmental conditions over time, it is expected that differential gene expression occurring throughout the growing season. Therefore, in a multi-harvest trial, a response variable can be treated as different traits in a multivariate framework analysis [2]. In this way, the genetic correlation between the same traits in different harvests is a measure of genotype by harvest interaction (G×H) [3-5]. Multi-harvest trials can also be described as a special case of multi-environment trials, in which the environments represent the different time points when data are collected in the same trial. The repeated measurement of the same trait over time generates longitudinal datasets and the sequential nature of measurements creates patterns of variation [6].

There are several models to deal with longitudinal datasets and the most common and simple (co)variance structure is the first-order autoregressive (AR1), where a single correlation parameter (*ρ*) is estimated. The model postulates a mechanism where the correlation between measurements *j* and *k* is *ρ*^|*j*―*k*|^, where the genotypic value of the genotype is a function of genes acting in a given time plus genes acting on the new measurement [3]. The AR1 model is an appealing method for modeling (co)variance structure for genotypes measured over time [3, 7, 8]. However, the AR1 model is recommended when the period between measurements is equally spaced, and when time points are unequally spaced nonlinear restrictions should be imposed for parameter estimation [9]. Due to the yield seasonality irregular time series are frequently observed in perennial forage yield measurements. The yield seasonality in forages is characterized by variation in growth and quality in response to environmental conditions [10]. Therefore, plants grow faster under favorable climate conditions and harvests are more frequent, whereas harvests are less frequent under unfavorable conditions. Thus, the time series for forage yield measures are naturally irregular.

Random regression models (RRM) were introduced by Henderson [11] and Laird and Ware [12]. Schaeffer and Dekkers [13] suggested their use in dairy cattle (*Bos taurus*) breeding to analyze day production records. Since then, several studies used RRM to predict growth in sheep (*Ovis aries*) [14], body weight in beef cattle [15], body weight in swine (*Sus scrofa domesticus*) [16], and egg production in layer (*Gallus gallus domesticus*) [17]. Recently RRM was applied to longitudinal data from perennial forage breeding trials for dry mater yield (DMY) in elephant grass (*Pennisetum purpureum* Schmach.) [18] and for forage quality in ryegrass (*Lolium perenne L*.) [19]. The use of RRM has also been increasing for annual crops with the advent of high-throughput phenotyping, which generates longitudinal datasets [20-22]. RRM can deal with longitudinal data (Schaeffer, 2004) as it captures the change of a trait continuously over time with few parameters by covariance functions (e.g., orthogonal polynomials and splines) [23, 24]. Kirkpatrick et al. [23] reported that RRM deals with unequally time-spaced measurements, relating that RRM should be the adequate model under this condition. Furthermore, there is a possibility to include environmental-dependent covariate in RRM, e.g., temperature and humidity to study the genotypes’ response to abiotic stress [25-27].

In this study, we investigated the use of RRM for longitudinal data of dry matter yield in ten forage breeding trials for three different species (alfalfa, guineagrass and brachiaria) for genotype selection and genetic interpretation.

## Material and methods

### Datasets

We used data from three forage species in 10 trials (T1 to T10) conducted from 2015 to 2020 in three locations and following different experimental designs (augmented row column design – ARCD, alpha-lattice design – ALD, and randomized complete block design – RCBD) (Table 1). Total dry matter yield (DMY, kg.ha^-1^) was assessed in all trials. The number of genotypes evaluated varied from eight (T4 and T5) to 182 (T1). The dataset is composed of data from early breeding trials (T1, T2, T3, T8, and T10) in which many genotypes are tested, and advanced breeding trials with a lower number of genotypes (T4, T5, T6, T7, and T9) (Table 1). The number of harvests in each trial varied from six (T7, T8, and T10) to 16 (T4 and T5).

**Table 1.**
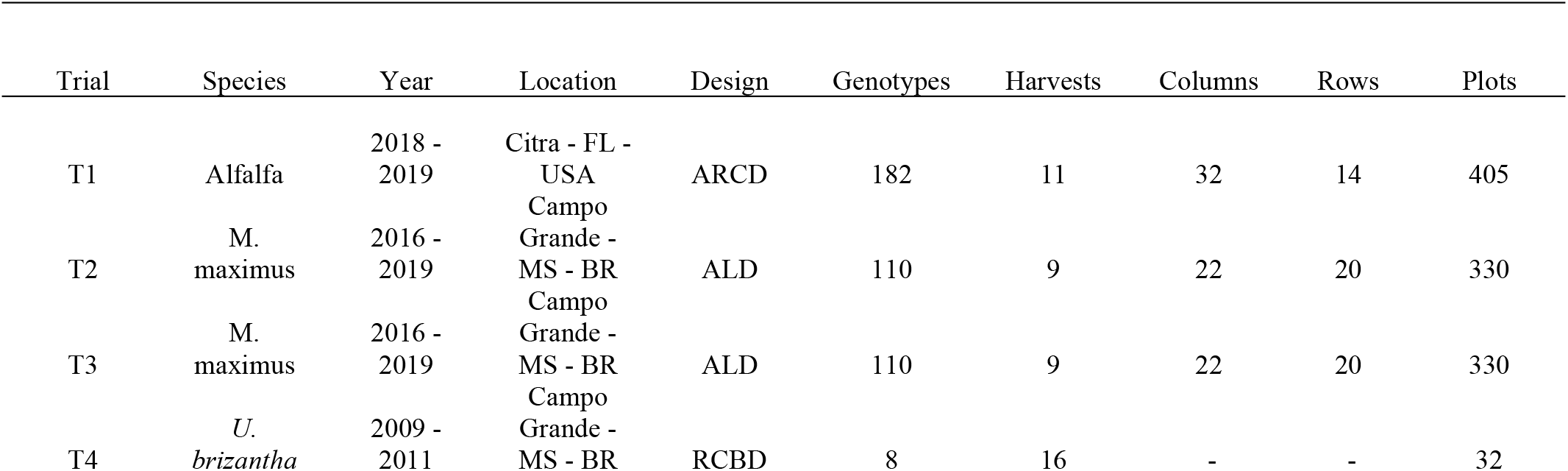
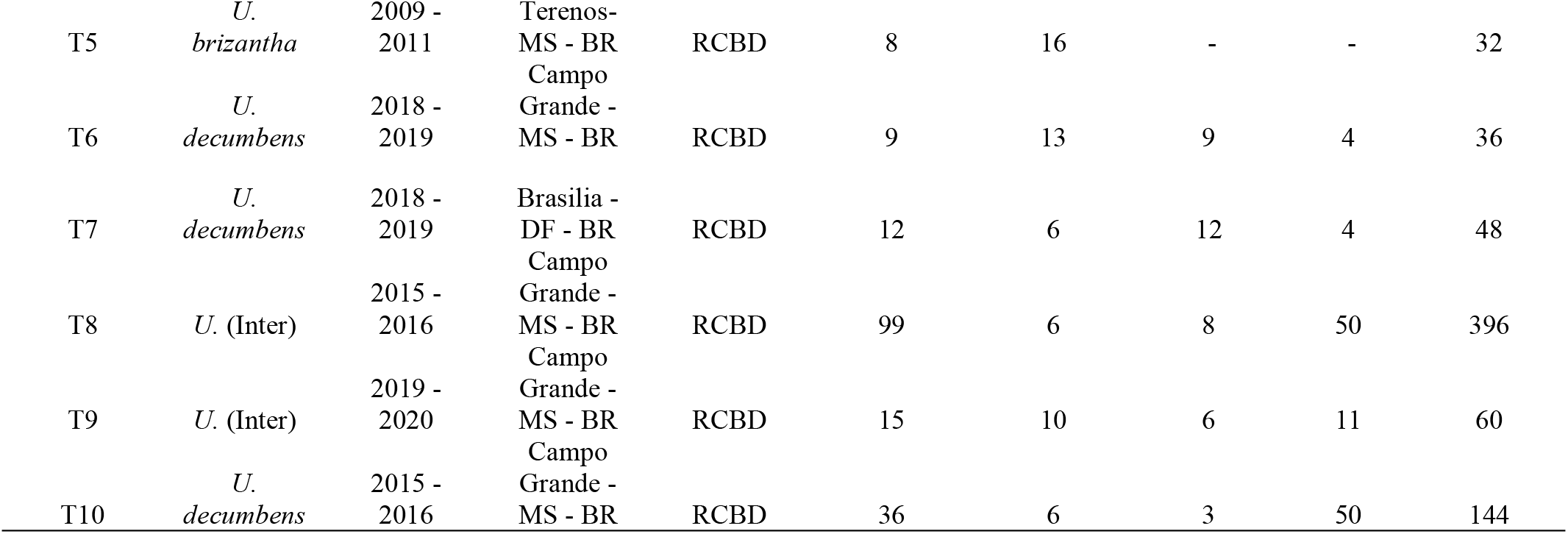
Description of experimental layout for the three forage species evaluated from 2015 to 2020.

### Statistical analysis

Analyzes for each multi-harvest trial (same trial over time) were performed based on a two-stage analysis using the weighting method proposed by Smith et al. [28]. In a two-stage analysis, genotypes’ best linear unbiased estimates (BLUEs) from each harvest in stage one were combined in a weighted multi-harvest mixed model analysis in stage two, where the weights provide a measure of relative uncertainty of the estimated genotypes’ BLUES from each harvest [28-30]. All the analyses were done by using ASREML-R [31] and SpATS [32] R packages, and the data summarization through graphs was done by ggplot2 [33] R package. The scripts for the analysis can be found at github (https://github.com/claudiocff/RRM-and-FAMM-asreml-two-step).

### First stage: estimating BLUEs and weights accounting for spatial variation

We obtained the BLUEs and weights of the genotypes at each harvest by trial, using the SpATS R package [32] in a mixed model framework:

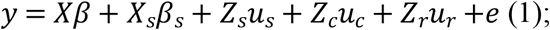

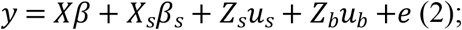

where, *y* is the vector from measured DMY from each plot; *β* is the vector of the fixed effects of genotypes and replication; *u*_*r*_ and *u*_*c*_ are the vectors of random effects of rows and columns, respectively in the augmented row column design (T1 - Table 1), where 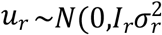 and 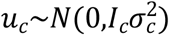; *u*_*b*_is the vector of random effects of block effects in a randomized complete block design (T4, T5, T6, T7, T8, T9 and T10 - Table 1), or the vector of the random effects of incomplete blocks within replication in the alpha-lattice design (T2 and T3 - Table 1), in which 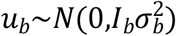 ; *β*_*s*_is the vector of fixed effects of the smooth spatial surface (unpenalized); *u*_*s*_is the vector of random effects of the penalized part of the smooth surface (penalized). The fixed term (*X*_*s*_*β*_*s*_- unpenalized) and random term (*Z*_*s*_*u*_*s*_- penalized) describe the mixed model expression of the smooth spatial surface (*f*(*r,c*) = *X*_*s*_*β*_*s*_ + *Z*_*s*_*u*_*s*_), where the random spatial vector *u*_*s*_has (co)variance matrix *S*. The SpATS model uses the P-spline ANOVA [PS-ANOVA, 34] to describe the 2D-splines in the mixed model framework. The *X*_*s*_, *Z*_*s*_incidence matrices, and the (co)variance matrix *S* are described by Lee et al [34] and Rodríguez-Álvarez et al. [32]. The PS-ANOVA parametrization can be decomposed as a linear sum of the univariate and bivariate smooth functions [35]; *e* is the vector of random errors, 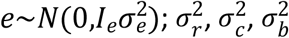 and 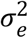are the variance components associated to the random effects of rows, columns, blocks and errors.*X, Z*_*r*_, *Z*_*c*_ and *Z*_*b*_are incidence matrices for the fixed effects, the random effects of rows, columns and blocks, respectively. *I*_*r*_, *I*_*c*_, *I*_*b*_, and *I*_*e*_are identity matrices.

### Second stage: modeling genotype by harvest interaction

For the statistical model described below, the BLUEs obtained for each genotype in each harvest by trial were regressed on a time gradient (days), where the first harvest of each trial were considered as day zero and the other harvest times were the days after the first harvest. Therefore, random coefficients were computed for each genotype to describe the ‘genotypes’ DMY trajectory over time. Polynomial functions were used to model the longitudinal dimensions by using orthogonal Legendre polynomials [23]. The orthogonal polynomials were obtained by rescaling the time points from -1 to 1 using the expression:

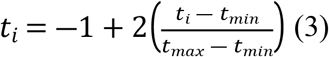

the Legendre polynomials are denoted by *P*_*n*_(*t*). Defining *P*_0_(*t*) = 1, the polynomial *n+1* is described by the recursive equation:

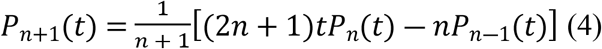

on the normalized form the Legendre polynomial can be described:

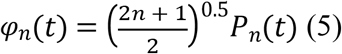

therefore, for a polynomial of order two we can obtain the following equations:

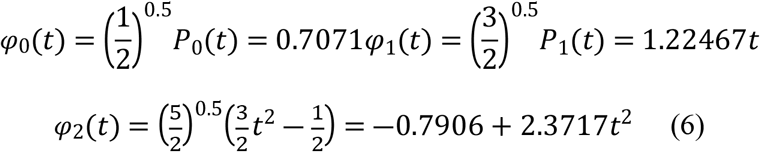

considering *m = 2* as the order of the covariance function to be used, the Legendre coefficient matrix *Λ* will have dimensions of (*m* + 1) × (*m* + 1) can be defined as:

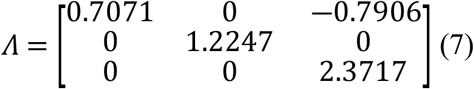

considering four different time points and an order two polynomial, the incidence matrix of time points (M) can be defined as:

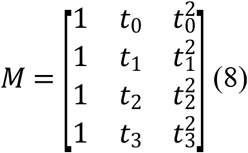

where *t*_*i*_ is the time point scaled by the equation (3).

Finally, the Legendre polynomials can be computed as *ϕ* = *MΛ*, where *ϕ* is a matrix containing the normalized polynomials for harvest time; *M* store polynomials of standardized harvest times; *Λ* is the matrix of Legendre polynomial coefficients of order *m+1*, where m is the degree of fit [36]. The random regression model can be defined as:

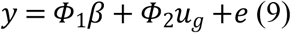

where *y* is the vector of BLUEs estimated by the models (1) or (2); *β* is the vector of the fixed regression coefficients; *u*_*g*_is the vector of random regression coefficients of genotypes, in which 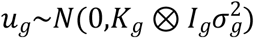; *e* is the vector of the errors, where 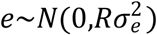. *K*_*g*_ is an unstructured (co)variance matrix associated to the random regression coefficients; the matrix *K*_*g*_ can be described as:

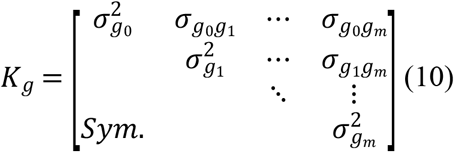

where, 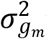 is the variance component associated to the coefficient of order *m*; *σ*_*g*__*n*_*g*_*m*_ is the covariance between the coefficient of order *m* and *n*.

*R* is the variance matrix of the errors, where *R* = *diag*(*R*_1_ *… R*_*j*_), in which *j* is the harvest by trial. In practice, *R*_*j*_ are unknown and replaced by an estimate 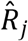 from each harvest. It is sometimes not feasible to store and use the full matrix 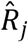 from each harvest, and so a vector of approximate weights is required. We used the weights proposed by Smith et al. [28], where the weights are based on the diagonal elements of 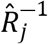 designated as *Π*, in which:

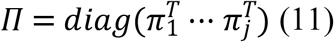

*π*_*j*_consists of the diagonal elements of 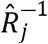. This simple approximation reflects the uncertainty in each estimated BLUE, accounting for within-trial heterogeneity, differing replication and spatial trends.

Based on Kirkpatrick et al. [23], the following estimator was used to obtain the genetic variance and covariance components across harvest times 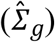 on original scale:

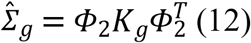

where, *ϕ*_2_is the incidence matrix of the Legendre polynomials associated to the random effects of genotypes; *K*_*g*_ is the (co)variance matrix associated to the random genotypes’ coefficients, defined in (10).

The genotypic values for each genotype across harvest time can be estimated by the equation:

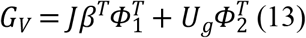

where *G*_*V*_ is a *i x j* matrix of the genotypic values on the original scale, where *i* is the number of genotypes and *j* the number of harvest time points; *J* is a column vector of 1’s size equal the number of genotypes (*i*); *β*^*T*^is the transposed vector of the fixed regression coefficients of size 1 *x* (*d+1)*, in which *d* is the degree of the polynomial fitted for the fixed regression; 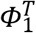 is the transposed incidence matrix of the Legendre polynomials for each harvest time for the fixed regression with size (*d+1) x j*; *U*_*g*_is the genotypes’s random coefficients matrix, size *i x* (m+1); 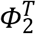 is the transposed incidence matrix of the Legendre polynomials for each harvest time for the random regression with size (*d+1) x j*.The polynomial function for the fixed regression was defined graphically by using a loess function, where the function order was determined by the number of curves (c) +1 in the mean DMY trajectory across harvest time (Fig 1). For example, for the trials T1 and T5 a polynomial of degree three were fitted, since two curves were observed on the mean DMY trajectory. The random polynomial regression degree was determined by the Bayesian information criteria (BIC).

**Fig 1.**
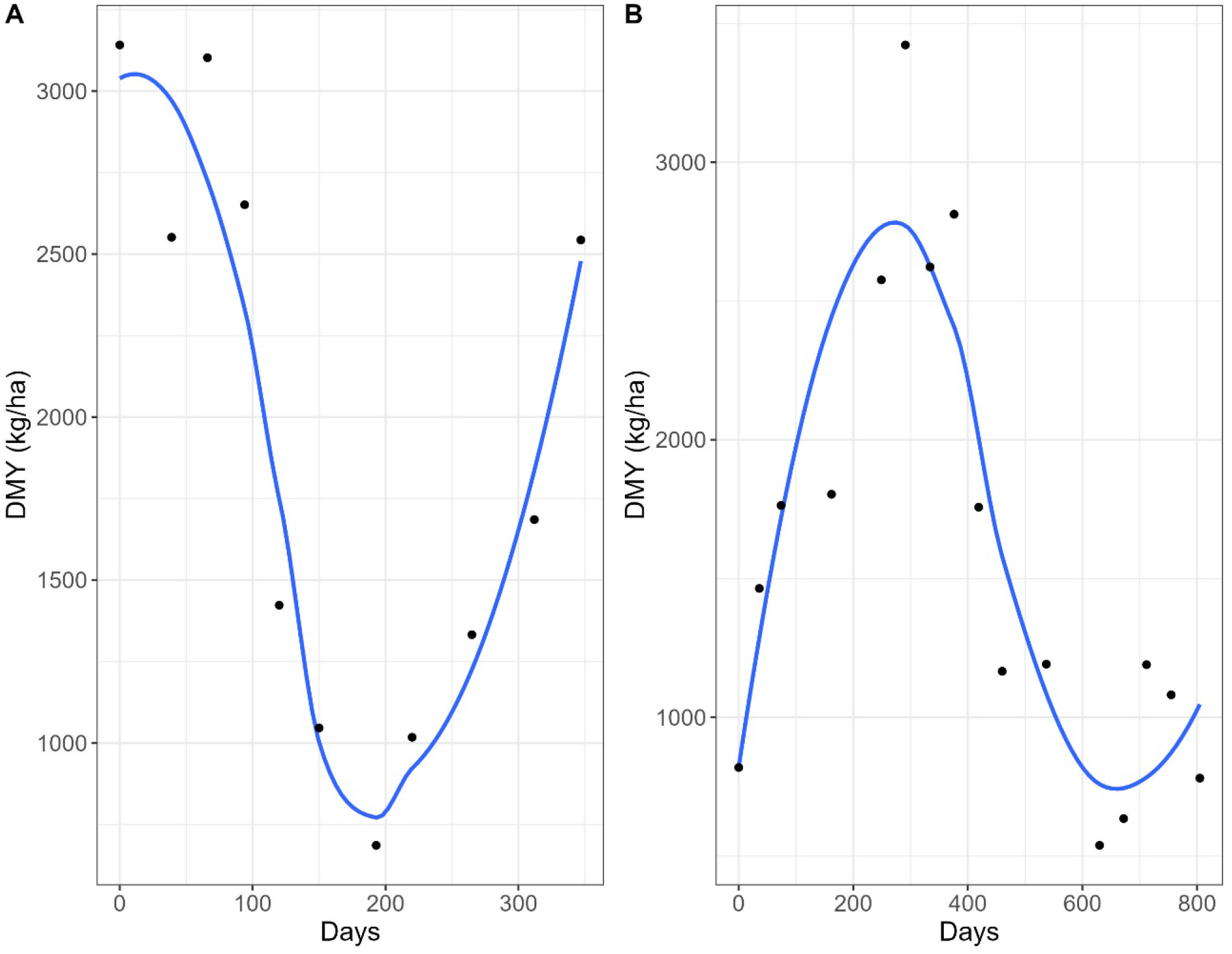
Mean dry matter yield (kg.ha^-1^) trajectory over time for trials T1 (Alfalfa – A) and T5 (Brachiaria – B)

### Heritability, broad adaptability, and stability

The heritability over harvest times was estimated by the expression:

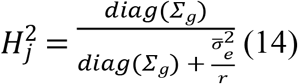

where 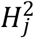 is the genotype mean-based heritability estimated at each harvest time; *Σ*_*g*_is the genetic variance-covariance matrix estimated by the equation (12); 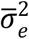is the mean error variance component across harvests; *r* is the number of replications in the trial.

The broad adaptability for each genotype was estimated based on the area under the DMY trajectory curve, in which reflects the total DMY accumulation over harvest time:

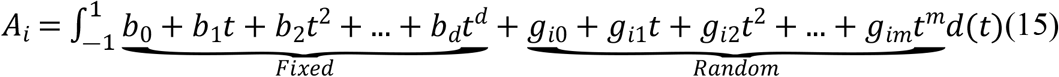

where *A*_*i*_ is the area under the trajectory curve of the genotype *i*; *b*_*0*_ is the fixed regression intercept; *g*_*i0*_ is the random regression intercept of the genotype *i*; *t* is the harvest time point; *d* and *m* are the polynomial fitted degree for the fixed and random regression, respectively; *b*_*d*_ is the fixed regression coefficient of degree *d; g*_*im*_is the random regression coefficient of degree *m* for genotype *i*.

The genotypes’ stability was calculated based on the trajectory curve’s coefficient of variation (*CV*_*c*_), in which reflect the genotypes’ Type I stability, where the genotype is stable if present small variance between environments (harvests), also called biological stability (Lin et al., 1986):

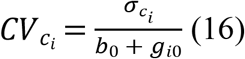

where,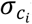is the standard trajectory curve deviation for genotype *i*; *b*_0_ + *g*_*i*0_is the overall performance for genotype *i*.

### Genetic interpretation on random regression models

One of the advantages of random regression models is the use of eigenfunction (*Ψ*_*k*_) of the genetic coefficient (co)variance matrix (10), in which can provide genetic insights about the studied trait based on Kirkpatrick et al. [23]:

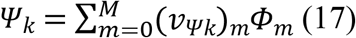

where (*v*_*Ψk*_)_*m*_is the *m*^*th*^ element of the *k*^*th*^ eigenvector of *K*_*g*_, and *ϕ*_*m*_ is the normalized value of the *m*^*th*^ Legendre polynomial.

## Results

### Overall description of RRM across trials

The degree of the polynomial used to fit the fixed part of the RRM varied from 2 (in T10) to 5 (in T2 and T3) (Table 2). However, there was no clear trend between the number of harvests evaluated in each trial and the order of the polynomial used (Table 1 and Table 2). The choice of polynomial degree for the fixed part of the model was based on the number of contrasting seasons evaluated in each trial. These contrasting seasons resulted in pits and peaks that had to be modeled by the fixed polynomial. On the other hand, for the random part of the RRM, lower-order polynomials were mostly preferred. In most trials, a first-order polynomial was sufficient to model the genotype by harvest interaction (G×H). A second-order polynomial was used in trials T4 and T8, and a third-order polynomial was employed in trial T1 (Table 2).

**Table 2.** Genetic parameters estimated using random regression models for alfalfa (T1), guineagrass (T2 and T3), and brachiaria (T4 to T10).

| Trial | Degree (F) | Degree (R) | $H^2$ | $\rho_g$ |
| --- | --- | --- | --- | --- |
| T1 | 3 | 3 | 0.36 (0.11) | 0.78 (0.19) |
| T2 | 5 | 1 | 0.63 (0.02) | 0.98 (0.02) |
| T3 | 5 | 1 | 0.51 (0.01) | 0.90 (0.10) |
| T4 | 3 | 2 | 0.34 (0.20) | 0.26 (0.54) |
| T5 | 3 | 1 | 0.52 (0.10) | 0.67 (0.33) |
| T6 | 3 | 1 | 0.16 (0.08) | 0.75 (0.28) |
| T7 | 4 | 1 | 0.42 (0.18) | 0.29 (0.61) |
| T8 | 4 | 2 | 0.69 (0.02) | 0.82 (0.18) |
| T9 | 3 | 1 | 0.31 (0.03) | 0.89 (0.11) |
| T10 | 2 | 1 | 0.55 (0.07) | 0.88 (0.11) |

Fitted polynomial order for fixed (F) and random (R) regression, mean heritability ( $H^2$ ) and genetic correlation between harvests ( $\rho_g$ ) for each trial estimated by RRM. The values between parenthesis represent the standard deviations across harvests for $H^2$ and among pair of harvests in case of $\rho_g$ .

Heritability varied across harvests, ranging from low (0.16 in T6) to median estimates (0.69 in T8) as shown in Table 2. Generally, there was little variation in heritability estimates across harvests, and high genetic correlations were observed between harvests, which suggests a higher predominance of non-crossover genotype by harvest interaction, except for trials T4 and T7 (Table 2).

The RRM models estimated an autoregressive covariance pattern for all datasets, indicating that closer harvests had higher correlations, while harvests that were further apart had smaller correlations (Fig 2). Trials T4, T5, T6, and T7 exhibited higher cross-over interactions, as evidenced by the negative correlations between harvests observed in these trials (Fig 2). The negative correlations could be attributed to the small number of genotypes evaluated in these trials (Table 1).

**Fig 2.**
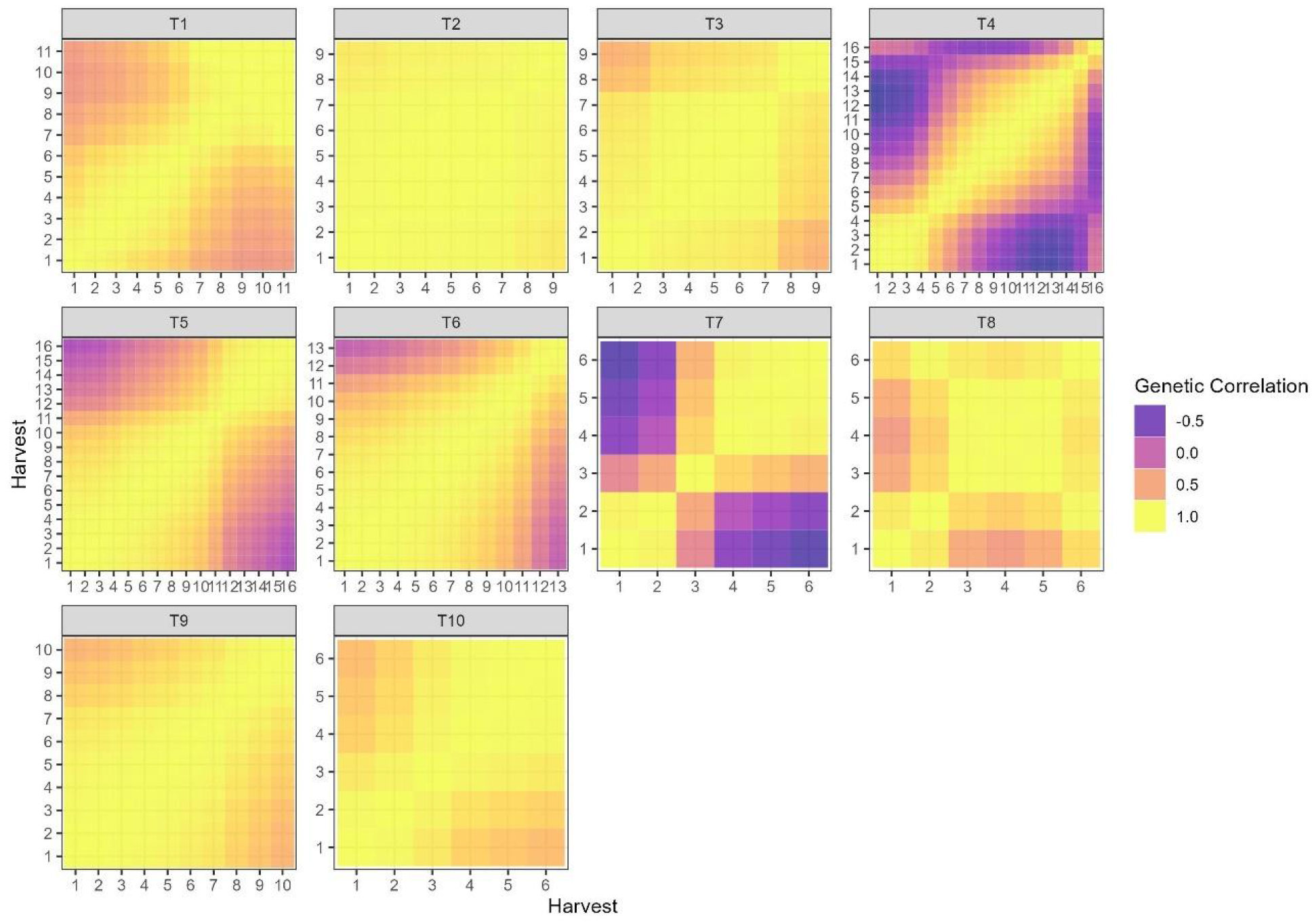
Heatmap of the genetic correlations between harvests estimated by RRM alfalfa (T1), guineagrass (T2 and T3), and brachiaria (T4 to T10).

In the upcoming sections of this study, we will provide an interpretation of the models for genotype selection. To ensure that the trials discussed have a high level of complexity in terms of genotype-environment interaction, we selected trials with low to median genetic correlation between harvests. Specifically, we chose trials T1, T5, and T7 for further interpretation (as shown in Table 2 and Fig 2).

### Genotype selection and G×H interpretation

#### Alfalfa breeding trial – T1

In this trial, 182 genotypes were evaluated over 11 harvests. To fit the model G×H for this trial, we used a Legendre polynomial of degree three for both the fixed and random effects (as shown in Table 2), which resulted in the estimation of 12 parameters.

### Variance components, heritability, and genetic behavior

The polynomial genetic variances varied from 2,930 to 343,654 for *g*_*3*_ and *g*_*0*_, respectively. The genotypes’ intercept (*g*_*0*_) retained most of the genetic variance, explaining 74% of the genetic variance, and the components related to the genotype’s curve shape (*g*_*1*_, *g*_*2*_ and *g*_*3*_) accounted to 26% of the genetic variance (Table 3). The most important correlation between the polynomial coefficients and the easiest to interpret is the correlation between *g*_*0*_ and *g*_*1*_, it shows the genetic variance behavior over time. In this trial, the correlation between *g*_*0*_ and *g*_*1*_ was negative (−0.22, Table 3), indicating lower genetic variance can be observed over time.

**Table 3.** Summary of genetic and non-genetic parameters estimated by RRM for alfalfa trial (T1).

| | $g_0$ | $g_1$ | $g_2$ | $g_3$ |
| --- | --- | --- | --- | --- |
| $g_0$ | 343,654 | -0.22 | 0.76 | 0.03 |
| $g_1$ | | 101,118 | -0.61 | -0.07 |
| $g_2$ | | | 19,627 | -0.63 |
| $g_3$ | | | | 2,930 |
| Importance (%) | 74 | 21 | 4 | 1 |
| $\bar{\sigma}_e^2$ | 365,436 | | | |

$g_0$ , $g_1$ , $g_2$ and $g_3$ are the random regression intercept and first, second and third order coefficient, respectively. $\bar{\sigma}_e^2$ is the mean error variance across harvests. The diagonal elements of the table are the genetic variances associated with the intercept and polynomial coefficients ( $\sigma_{g_0}^2, \sigma_{g_1}^2, \sigma_{g_2}^2, \sigma_{g_3}^2$ ); the off diagonal of the table are the correlations between intercept and polynomial coefficients ( $\rho_{g_0g_1}, \rho_{g_0g_2}, \rho_{g_0g_3}, \rho_{g_1g_2}, \rho_{g_1g_3}, \rho_{g_2g_3}$ ).

One advantage of RRM is the heritability estimation in function of time, as well as pointed out above by the negative estimate of *ρ*_*g*0*g*1_the heritability tended to decrease over time as the genetic variance also decreased. The lower heritability estimates (*H*_*2*_ < 0.30) ocurred between 120 and 220 days after the first harvest (Fig 3). It is noteworthy that the harvest with lower heritability occurred during the late summer to late fall period when alfalfa genotypes are exposed to various stress factors such as high temperatures, cloudy days, and high humidity, along with biotic stress mainly caused by fungal diseases (see Fig. 3).

**Fig 3.**
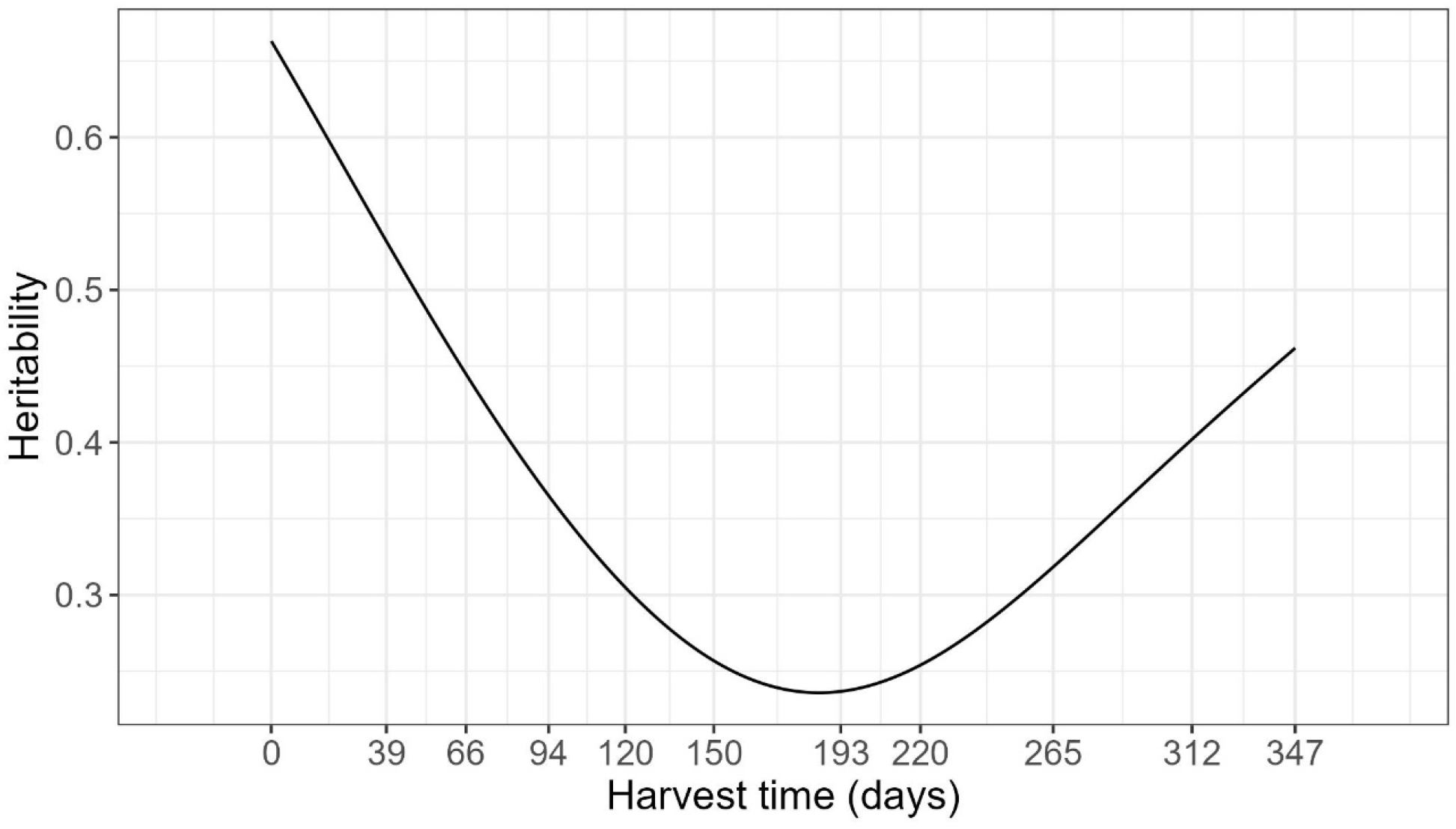
Heritability trajectory over harvests time (days) for alfalfa trial (T1).

The genetic correlation varied from 0.43 to 1.00 (Fig 4A). The genetic correlation between harvests followed an autoregressive pattern, where harvest closer to each other tend to have higher genetic correlation and those harvests far apart from each other had lower genetic correlation (Fig 4A). The eigen functions can be used in RRM to infer about gene expression over time (Fig 4B). Where the first eigenfunction had a nearly constant behavior and explained 74% of the genetic variation, this variation represents a common gene poll that is being expressed over time, and explain the simple G×H interaction, since non differential expression was observed (Fig 4B). The second eigenfunction represents another gene pool, in which explained 21% of the genetic variation that shows differences in gene expression under different environment conditions, explaining most of the complex G×H interaction (Fig 4B). The third and fourth eigenfunctions explained only 4 and 1% of the genetic variation, also representing complex G×H interactions, where differences on gene expression can be observed over time (Figure 4B).

**Fig 4.**
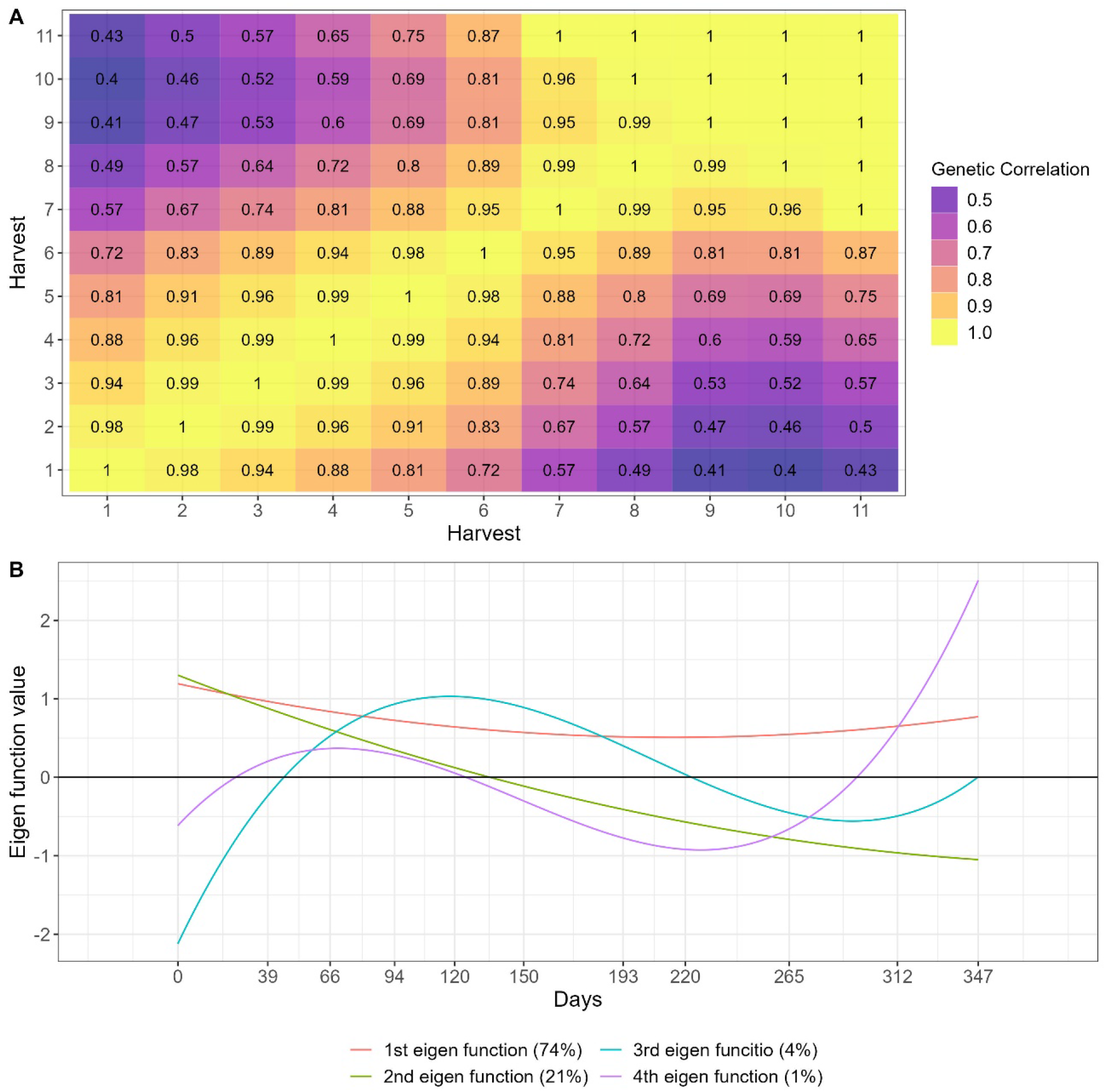
Heat map (A) and Eigenfunction smooth curves (B) illustrating genetic correlations between alfalfa harvests (T1).

### Genotypes’ adaptability, stability, and yield trajectory over time

Forage breeders are interest in the genotype’s behavior over time, a good genotype should have higher yield over time. Therefore, RRM can be a useful tool where genotypes’ reaction norms can be plotted (Fig 5). There was great variability on the genotypes’ reaction norm, and the main changes on ranking occurred between harvests performed between 39 to 220 days. On Fig 5, we highlighted seven genotypes that presented higher area under the curve (*A*), i.e., higher broad adaptability, the trial’s checks (UF2015, FL99 and B_805), and the mean yield trajectory curve. It can be inferred from Fig 5 that genotypes 15F, 103F, 33_H, and 42F exhibited superior performance during the stress period (from day 150 to day 265) and can therefore be considered as more tolerant genotypes. All seven genotypes with higher *A* performed better than the checks for most of the harvest evaluated (Fig 5).

**Fig 5.**
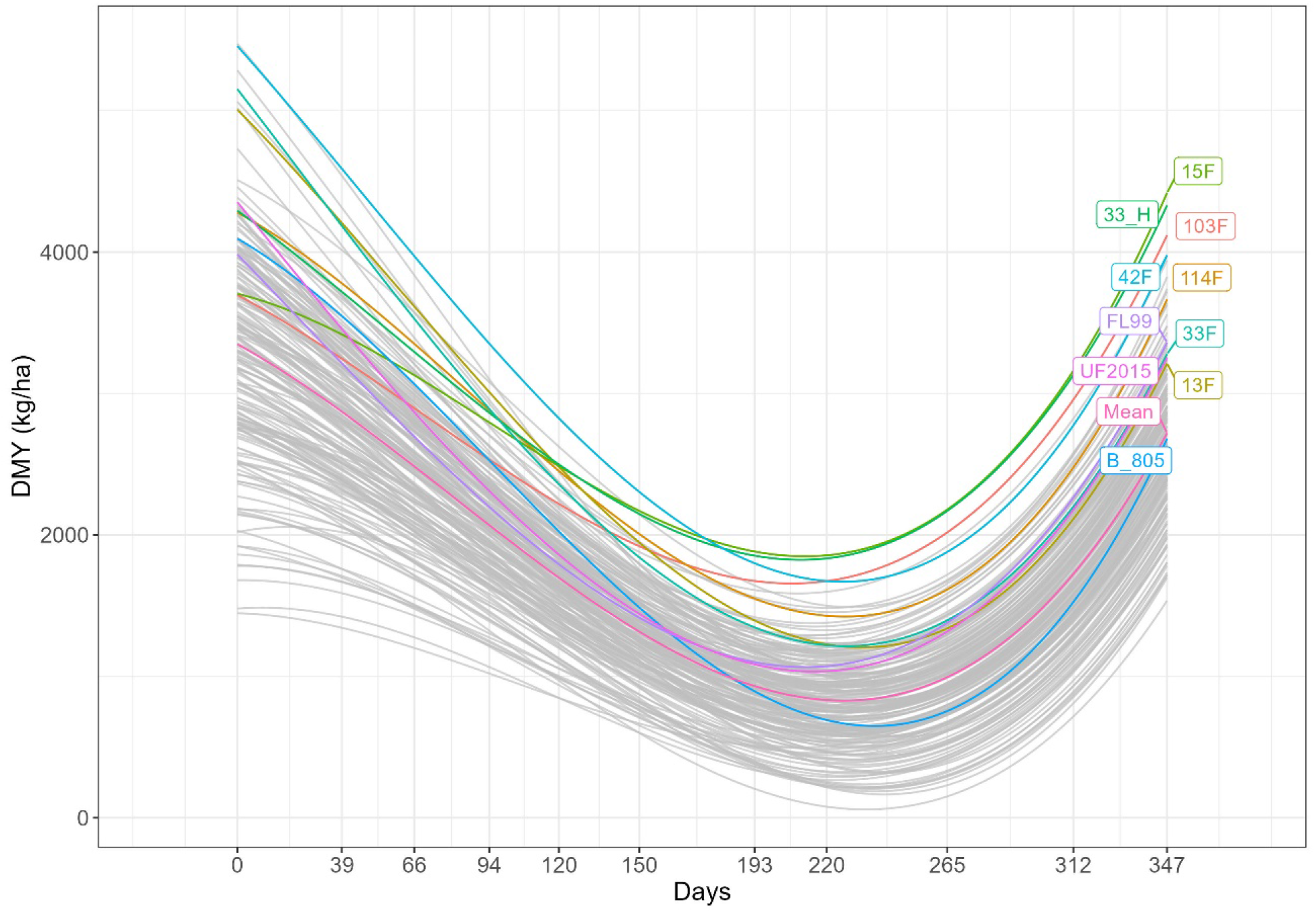
Genotypes’ dry matter yield (kg.ha^-1^) trajectory over harvest time for alfalfa trial (T1). The Highlighted genotypes represents the better genotypes based on the area under the curve, the checks (B_805, UF2015 and FL99) and the mean DMY trajectory.

Although, genotype selection based only on RRM is not feasible when large breeding populations are evaluated and complex G×H interactions are significant. To overcome this difficulty, we proposed the genotype selection based on the area under the curve (*A)* and in the curve coefficient of variation (*CV*_*c*_), which represent genotypes’ adaptability and stability for DMY, respectively. It was observed high variability for *A* and *CV*_*c*_ between genotypes (Figs. 6A and 6B), and the correlation between the two parameters was of median magnitude (−0.55, Fig. 6A), thus it is possible to select genotypes with higher *A* and lower *CV*_*c*_, i.e., genotypes presenting high adaptability and stability. For selection purposes we did a scatter plot showing the genotype’s *A* and *CVc* values, where the genotypes to be selected are on the superior left quadrant of the graph (Fig. 6A). The 10% genotypes with higher *A* are highlighted on Fig. 6A.

**Fig 6.**
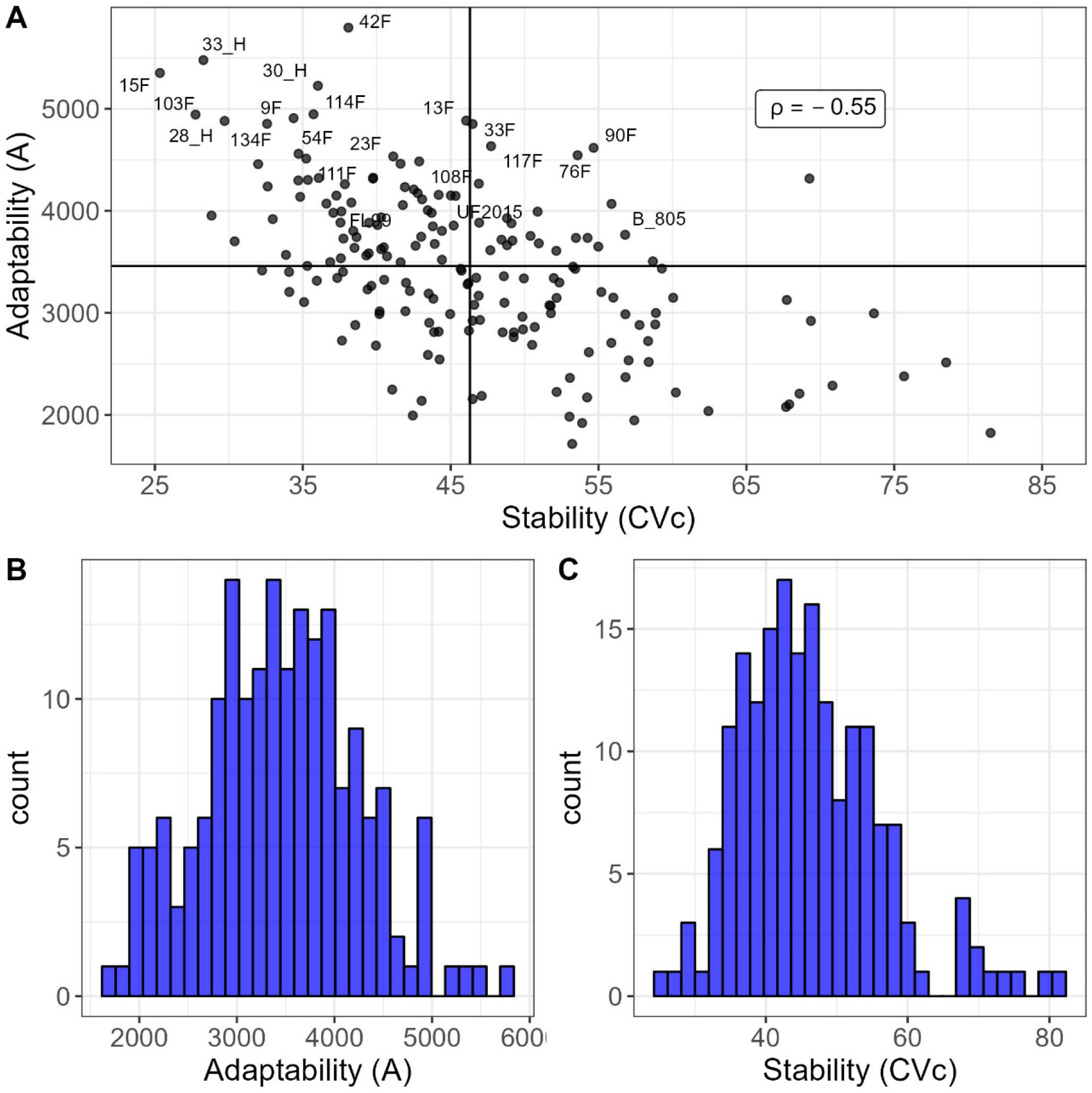
Scatter plot of Stability versus Adaptability (A) with solid black lines indicating mean values of *A* and *CV*_*c*_ for the alfalfa trial (T1); Histogram of Adaptability (B); Histogram of Stability (C).

### *Urochola brizanta* advanced breeding trial – T5

In this trial, 9 genotypes were evaluated for DMY across 16 harvests. The degree of Legendre polynomial fitted for this trial was three for fixed and one for random part of the model (Table 2). Seven parameters were estimated for RRM.

### Variance components, heritability, and genetic behavior

For this trial only the intercept and first order polynomial coefficient were needed to model the G×H interaction. The intercept (*g*_*0*_) and slope (*g*_*1*_) explained 69 and 31% of the genetic variance (Table 4). The negative correlation (*ρ*_*g*0*g*1_ = ―0.41) between *g*_*1*_ and *g*_*0*_ indicated that genetic variances decreased over time. The highest heritability estimates occurred between harvests performed between 0 to 71 days (*H*_*2*_ > 0.70), and the lowest estimates between 450 and 619 days (*H*_*2*_< 0.43) (Fig 7). The first drought season (71 to 238 days – May to October/2009) had heritability estimates varying from 0.57 to 0.69, whereas the second drought season (450 to 619 – May to October/2010) presented the lower heritabilities varying from 0.40 to 0.43 (Fig 7).

**Table 4.**
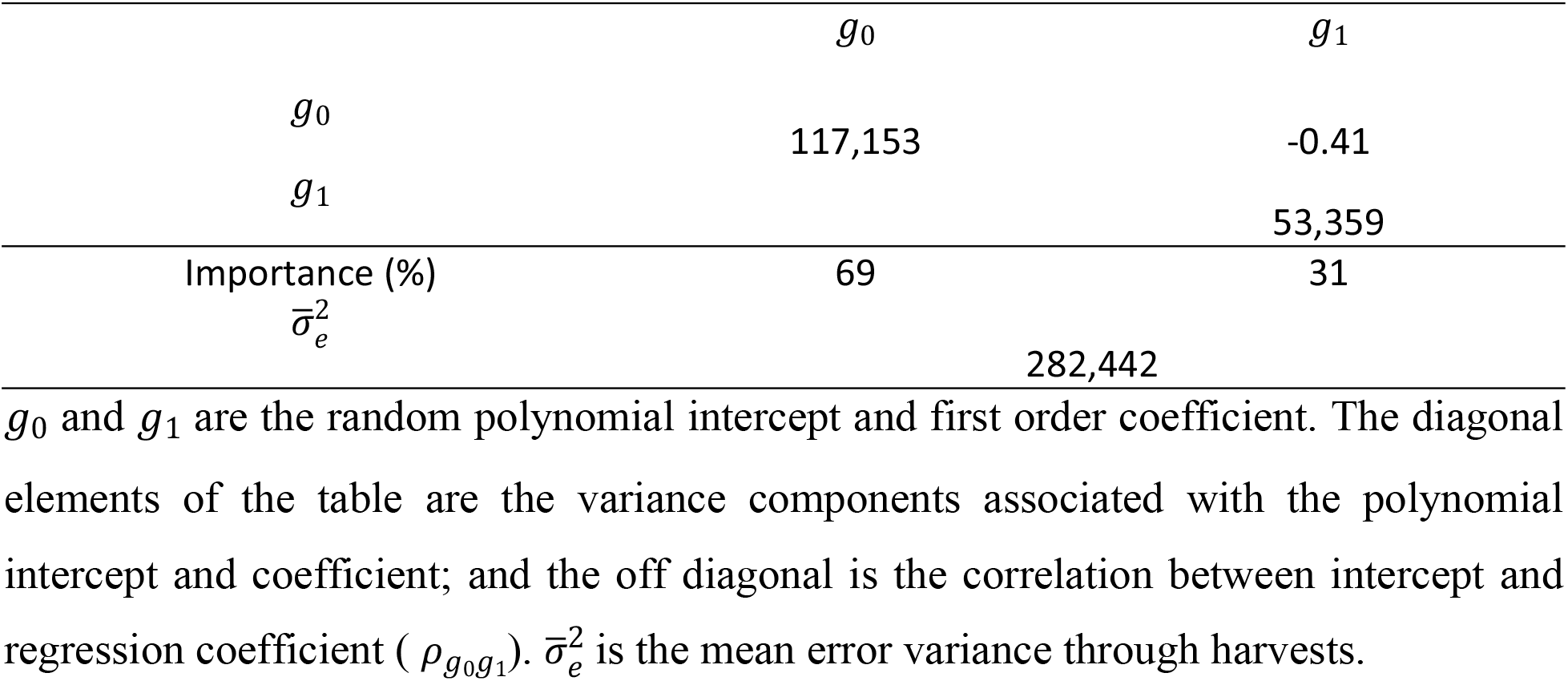
Summary of genetic and non-genetic parameters estimated by RRM for trial T5.

**Fig 7.**
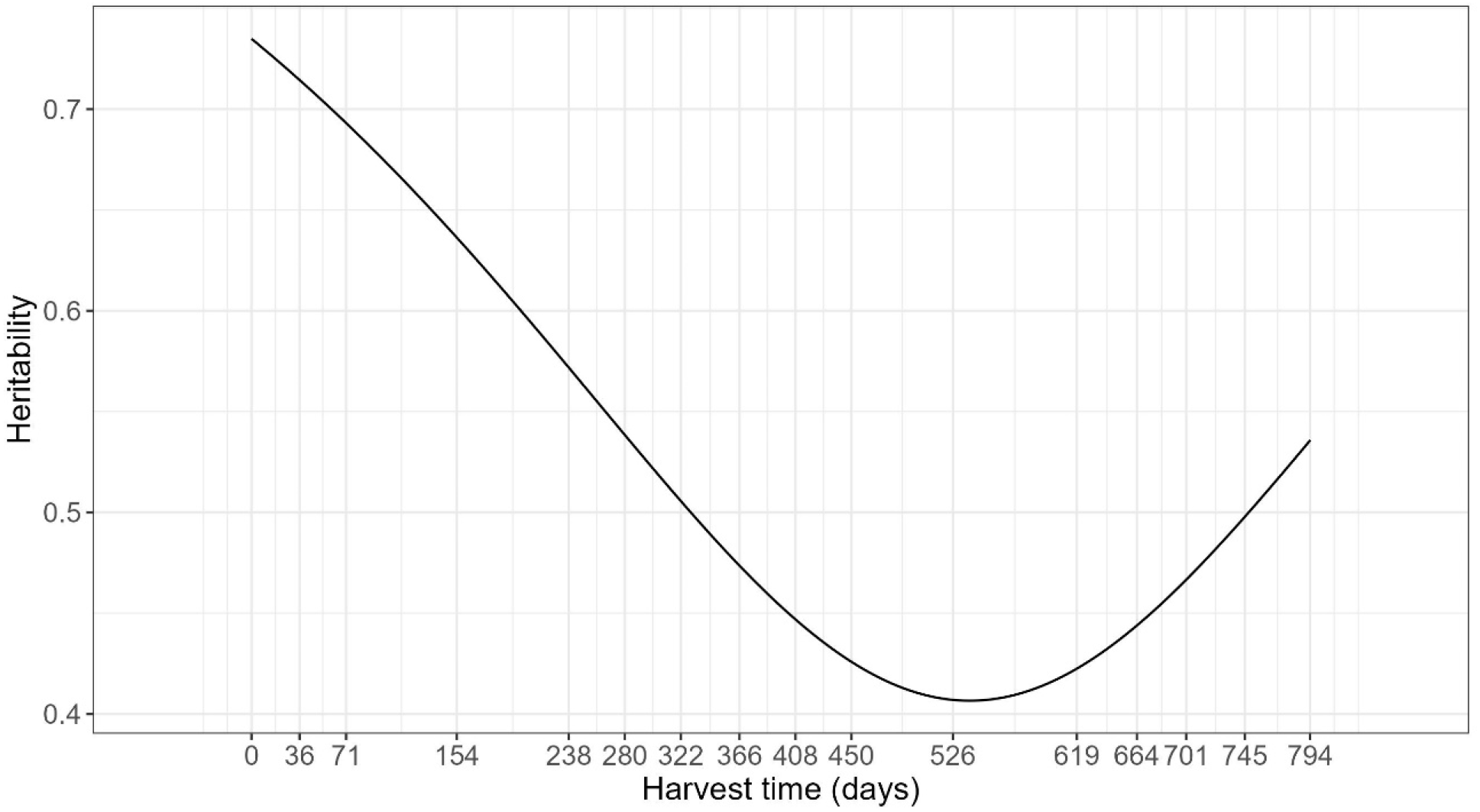
Heritability trajectory over harvests time (days) for brachiaria trial (T5).

The genetic correlation across harvests varied from -0.17 to 1.00 (Fig. 9A). Like T1 (Fig 4A), the genetic correlations on T5 followed an autoregressive structure (Fig 8A), but there was no common factor explaining the G×H interaction for all harvests as negative genetic correlations occurred in this trial (Fig 8A). This fact can also be explained by the eigenfunctions, where the two eigenfunctions varied over time (Fig 8B). First and second eigenfunctions explained the complex G×H interaction, where the gene expression varied over time for the two eigenfunction (Fig 8B).

**Fig 8.**
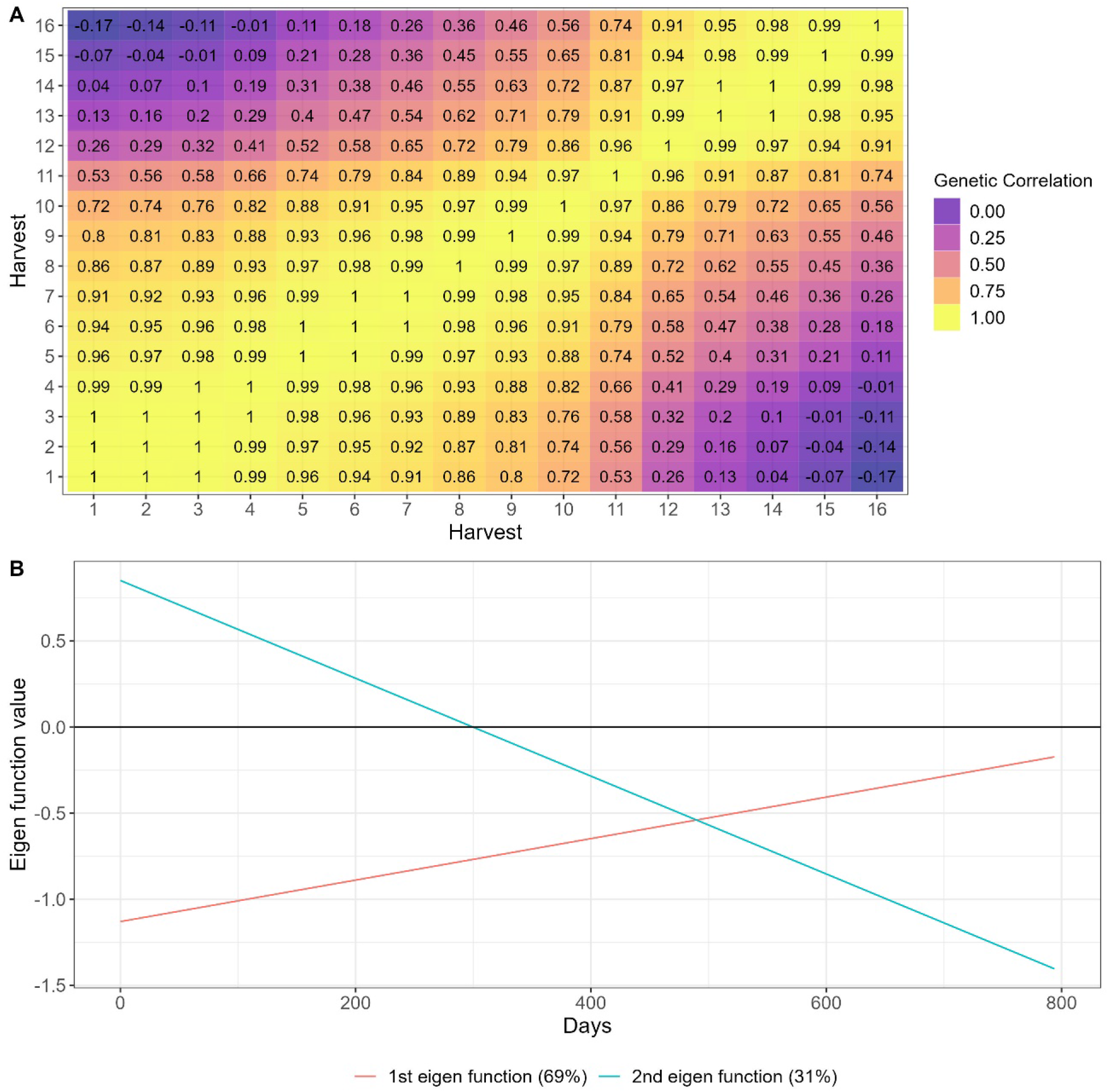
Heat map (A) and Eigenfunction smooth curves (B) illustrating genetic correlations between brachiaria harvests (T5).

**Fig 9.**
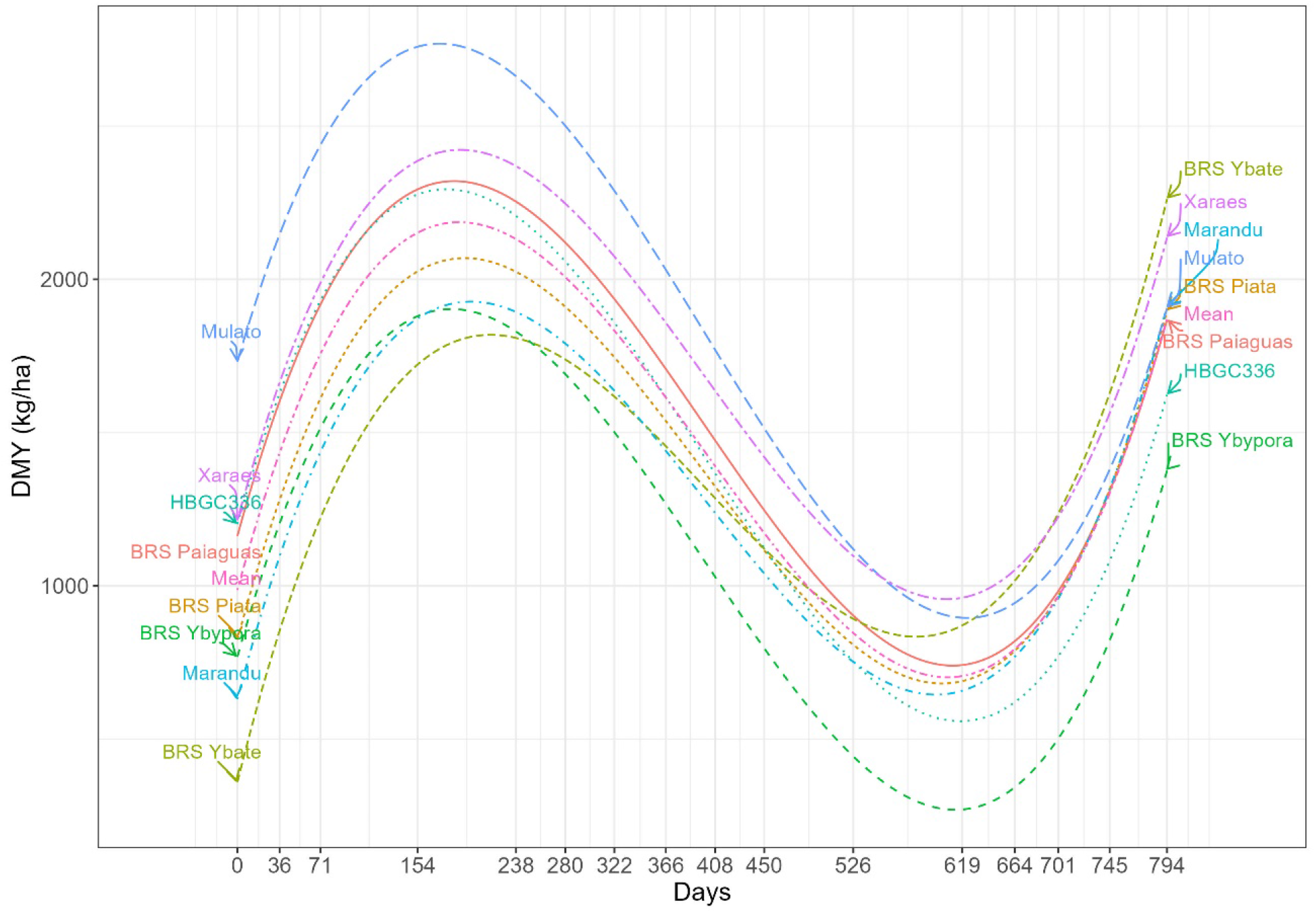
Genotypes’ dry matter yield (kg.ha^-1^) trajectory over harvest time (days) for trial T5.

### Genotypes’ adaptability, stability, and yield trajectory over time

The genotypes’ reaction norms (Fig 9) showed that most changes in genotypes raking occurred between days 238 and 664. The maximum DMY was reached between the first drought season (154 days) and beginning of the second rainy season (238 days) (Fig 9). This is an atypical behavior and can be explained by the time interval between harvests, reflecting on a longer period of dry matter accumulation (Fig 9). Another factor is the number of harvests realized before this period where only three harvests were done until the first drought season (Fig 9). Furthermore, atypical climate conditions could happen in this season. As expected, the lowest DMY occurred at the end of the second drought season (619 days) (Fig 9). By the atypical behavior of the genotypes on first drought season, the selection of tolerant genotypes to this condition should be done by looking at genotypic values in the period from 450 to 619 days (Fig 9). Under drought conditions the genotypes BRS Ybapé, Xaraés and Mulato had the greatest DMY (Fig 9). However, BRS YBAPÉ had the lowest DMY at the first harvest, perhaps due to a poor establishment (Fig 9). By the reaction norms the genotype Mulato had the best performance across harvests with good establishment as well as good performance under the drought season (Fig 9). The correlation between adaptability and stability was -0.54, indicating that selection can be done for both parameters (Fig 10). Three genotypes can be selected by presenting higher stability and adaptability (Mulato, Xaraés and BRS paiaguás, Fig 10).

**Fig 10.**
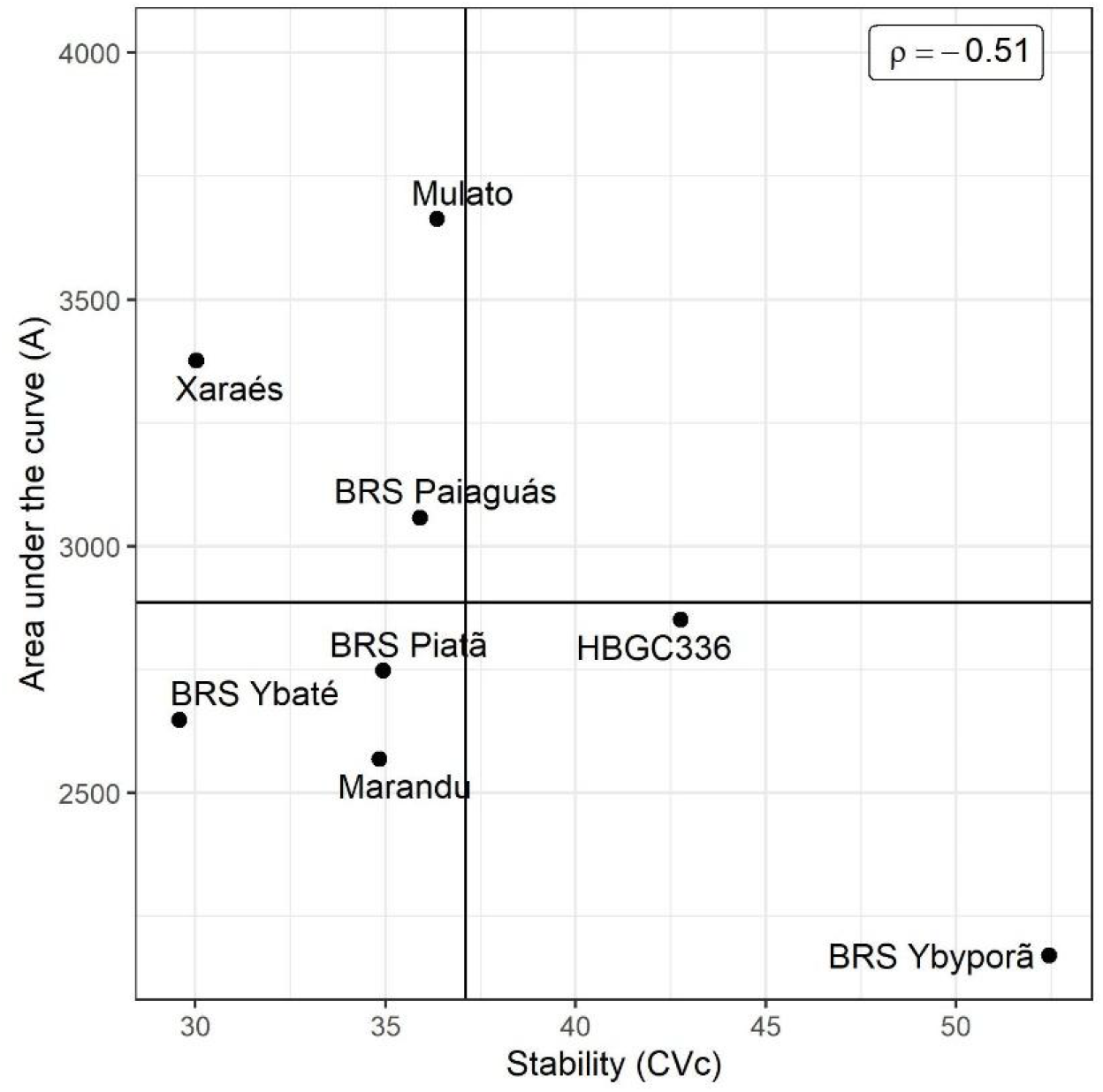
Scatter plot depicting the relationship between genotypes’ stability (measured by coefficient of variation - *CV*_*c*_) and adaptability (measured by area under the curve - *A*)

### *Urochola decumbens* advanced breeding trial – T7

In this trial, 12 genotypes were evaluated for DMY across six harvests. The degree of Legendre polynomial fitted for this trial was four for fixed and one for random part of the model (Table 2).

### Variance components, heritability, and genetic behavior

The intercept explained a small amount (35%) of genetic variance than the first order polynomial coefficient in which explained 65% of the genetic variance (Table 5). Furthermore, the correlation between slope and intercept was positive, indicating that genetic variance increased over time (Table 5). The heritability estimates varied from 0.20 at the third harvest (drought season – August/2018) to 0.82 at the last harvest (beginning of drought season – June/2019) (Fig 11).

**Table 5.**
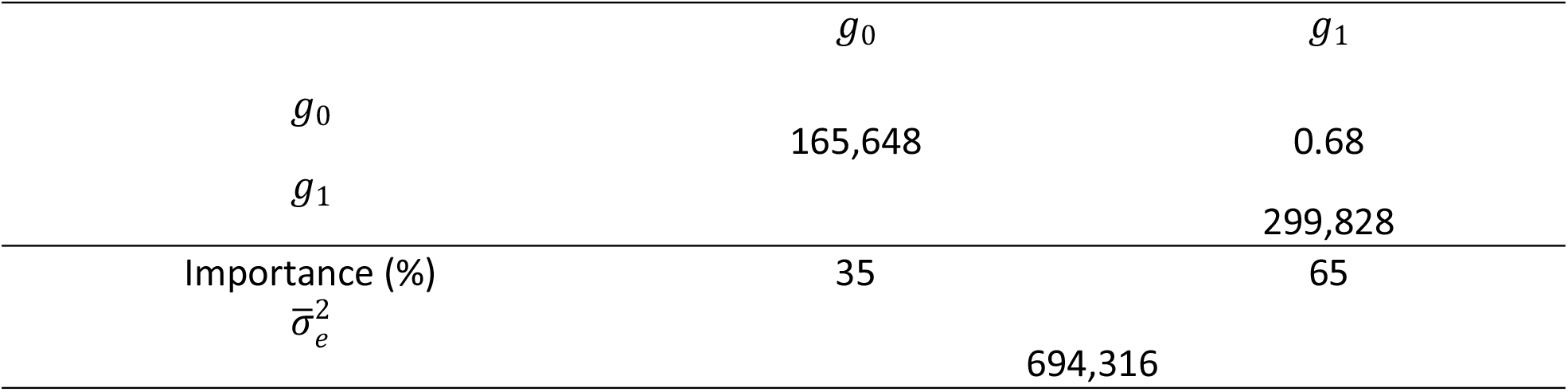
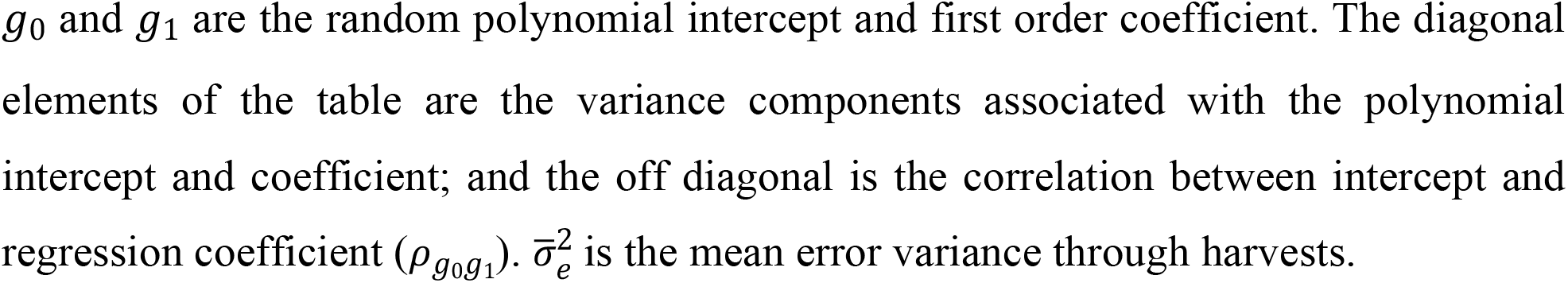
Summary of genetic and non-genetic parameters estimated by RRM for trial T7.

| | $g_0$ | $g_1$ |
| --- | --- | --- |
| $g_0$ | 165,648 | 0.68 |
| $g_1$ | | 299,828 |
| Importance (%) | 35 | 65 |
| $\bar{\sigma}_e^2$ | 694,316 | |

$g_0$ and $g_1$ are the random polynomial intercept and first order coefficient. The diagonal elements of the table are the variance components associated with the polynomial intercept and coefficient; and the off diagonal is the correlation between intercept and regression coefficient ( $\rho_{g_0g_1}$ ). $\bar{\sigma}_e^2$ is the mean error variance through harvests.

**Fig 11.**
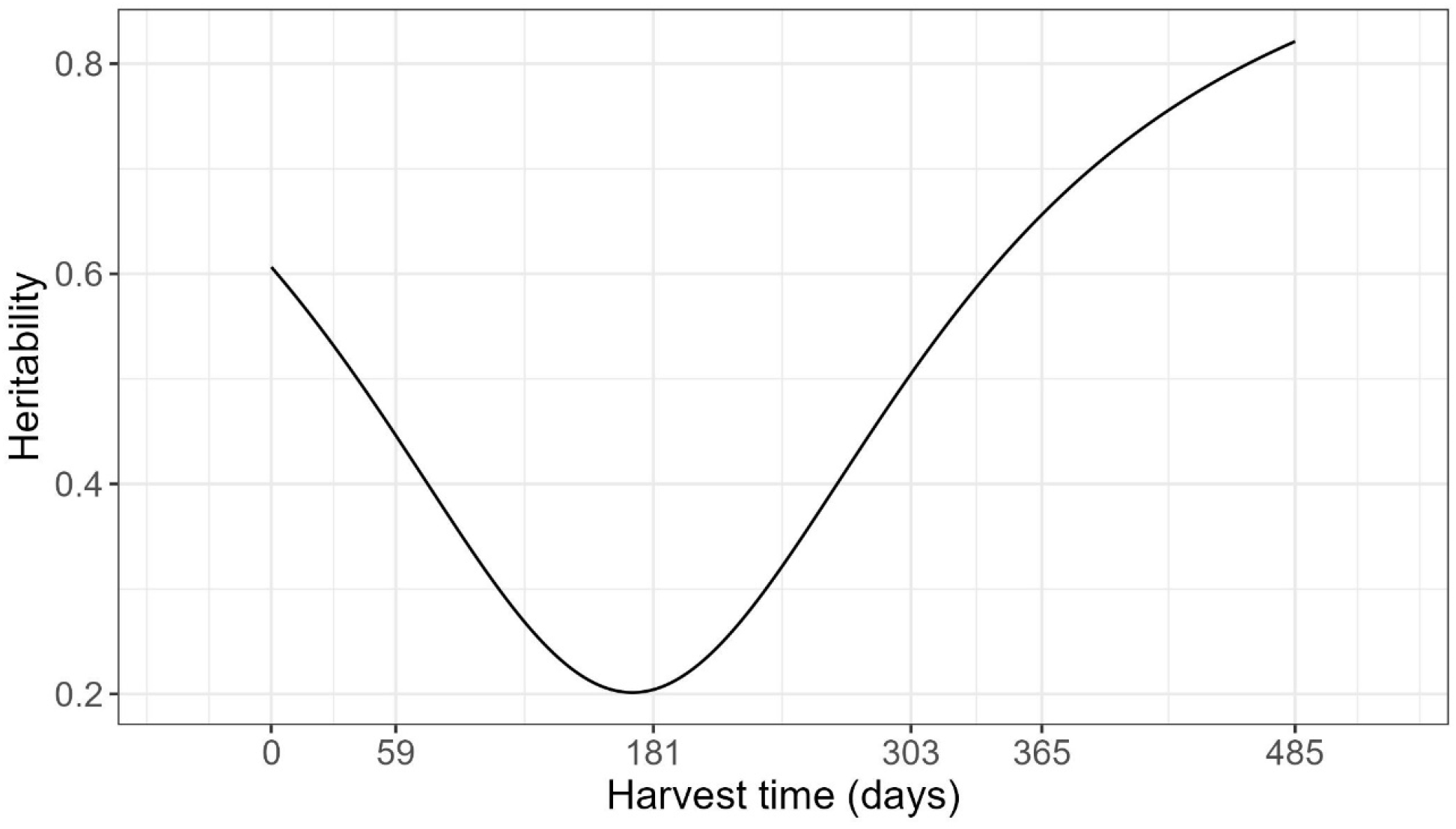
Heritability trajectory over harvests time (days) for brachiaria trial (T7).

The genetic correlations across harvests varied from 0.99 to -0.79, indicating a high and complex G×H interaction effects (Fig 12A). As it happen for T1 and T5, the RRM approximated the (co)variance structure into an autoregressive structure on T7 (Fig 12A). The two estimated eigenfunctions varied across time, indicating there was not a common gene pool expressing in the same way for all harvests evaluated (Fig 12B). Both gene pools represented by the eigenfunctions are expressing differentially across harvest, explaining the strong complex G×H interaction occurred for this trial (Fig 12B).

**Fig 12.**
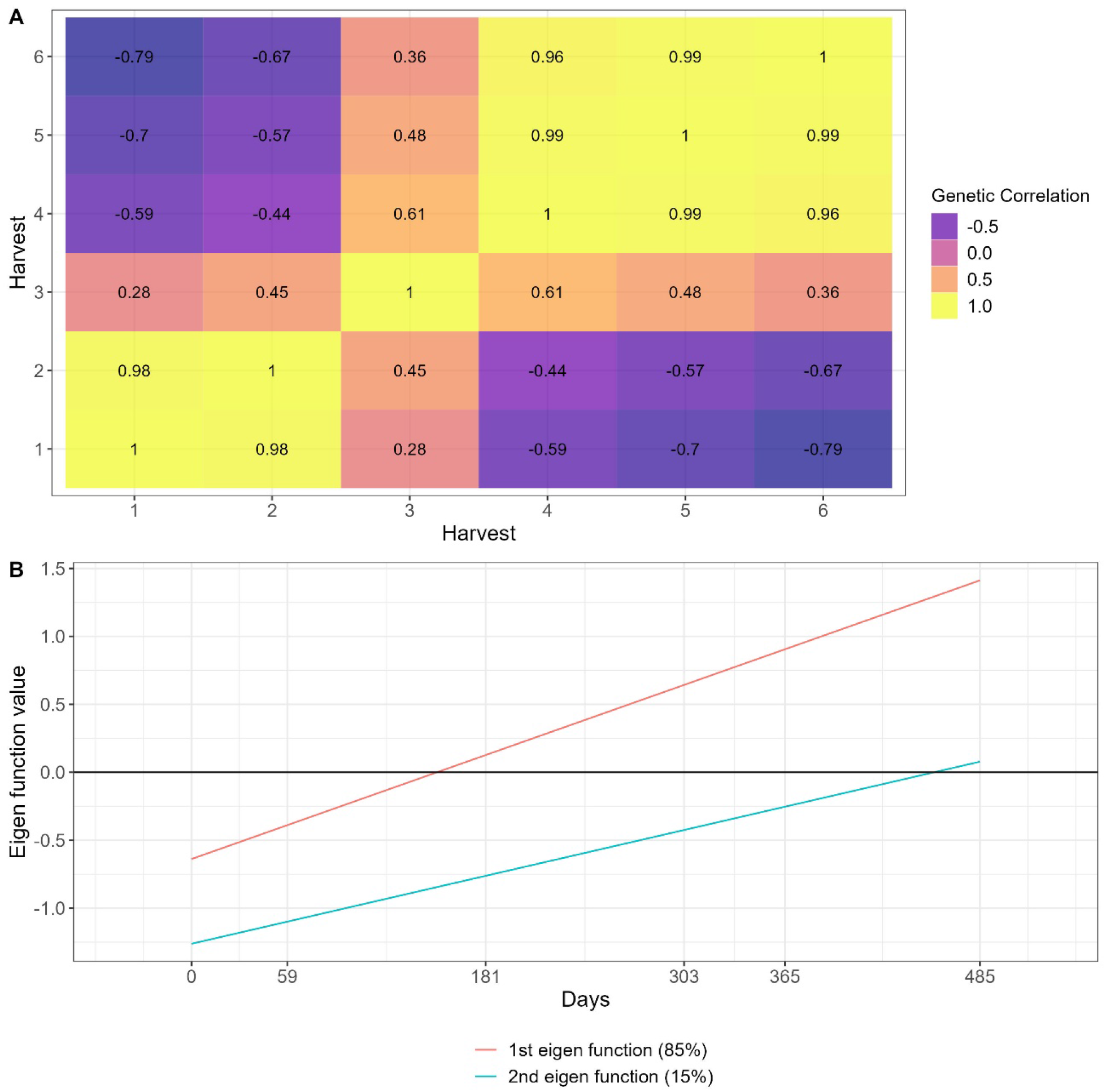
Heat map (A) and Eigenfunction smooth curves (B) illustrating genetic correlations between brachiaria harvests (T7).

### Genotypes’ adaptability, stability, and yield trajectory over time

The most changes in genotypes raking occurred after the first drought season at harvest performed on day 181 (Fig 13). The highest DMY occurred at day 365 (rainy season), and the lowest DMY occurred at the first drought season (day 181) (Fig 13). The genotype BRS YBATÉ, also evaluated in trial T5, had the same behavior, where it had a poor establishment (day 0 to 59) and a good recovery after the first drought season, being the best genotype in all harvests after the drought season (Fig 13). The correlation between *CV*_*c*_ and *A* was -0.25, indicating the possibility of selecting adaptable and stable genotypes (Fig 14). Although the genotype BRS YBATÉ had the best adaptability, it was one of the most unstable genotypes with greater variation in DMY across harvests (Fig 14). Five genotypes were identified having good stability and adaptability (HD1, HD3, Basilisk, Paiaguás and HD4) (Fig 14).

**Fig 13.**
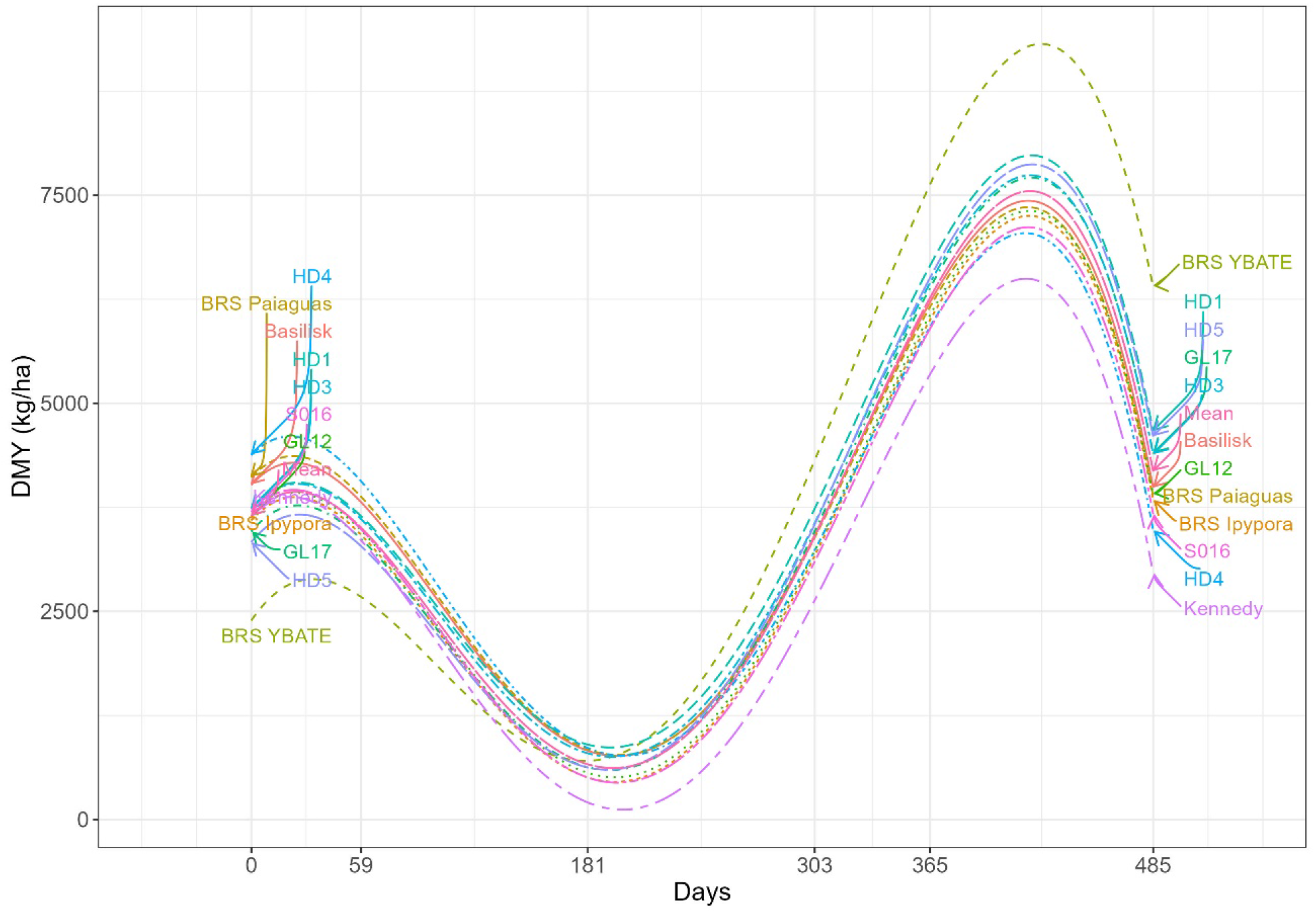
Genotypes’ dry matter yield (kg.ha-1) trajectory over harvest time (days) for trial T7.

**Fig 14.**
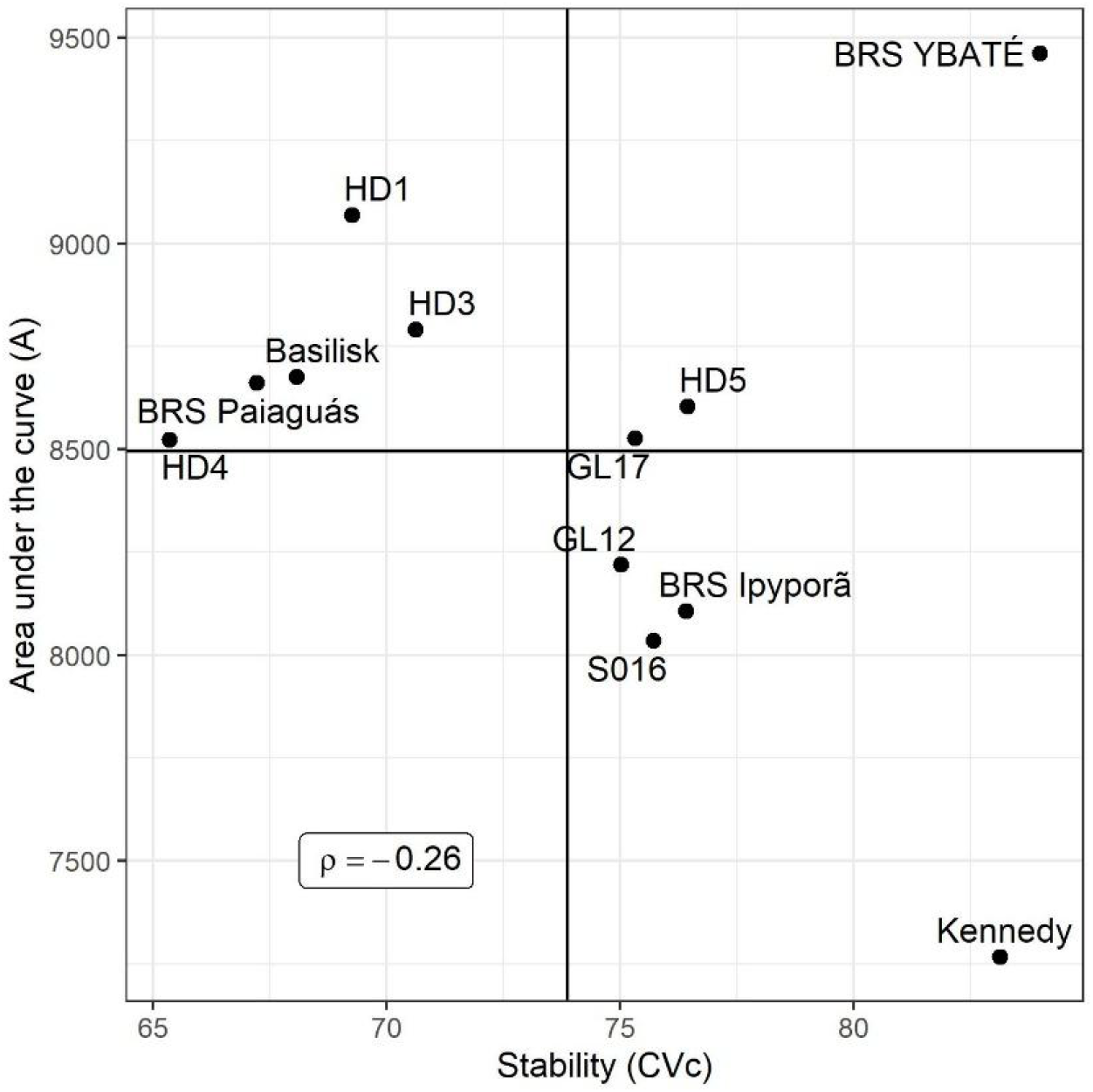
Scatter plot depicting the relationship between genotypes’ stability (measured by coefficient of variation - *CV*_*c*_) and adaptability (measured by area under the curve - *A*)

## Discussion

Our study focused on the analysis of longitudinal DMY data from ten different trials and four perennial forage species, utilizing RRM methodology. Our approach included the estimation of variance components and the selection of genotypes based on their adaptability and stability for DMY. Through this analysis, we aim to gain deeper insights into the performance of different genotypes over time, and to identify the most suitable candidates for future breeding programs.

### Goodness of fit evaluation

Proper modeling of genetic effects over time is crucial for accurate analysis of yield data in forage perennial species [37]. RRM has proven to be an effective method for dealing with longitudinal records in animal breeding [38, 39] and can be adapted for use in analyzing longitudinal data in plants. As observed in all trials presented in this study, DMY records were obtained at unequally spaced intervals due to the seasonality inherent in forage growth under tropical and subtropical climates [10]. Infinite-dimensional methods, such as RRM, offer an advantage in this context, as they can effectively handle unequally spaced records. This is because yield trajectories are continuous functions of time, meaning that an infinite rather than finite number of measurements is required to fully describe a trait in an individual [23,40].

The selection of the random polynomial order in RRM can be achieved by using goodness of fit and parsimony criteria, as demonstrated by Corrales et al. [41]. In this study, we adopted the Bayesian information criterion (BIC) approach, as suggested by Rocha et al. [18], to select the random polynomial order. However, choosing an appropriate fixed function to model the phenotypic curve shape can be challenging. In cases where the overall trajectory is linear, fitting the fixed part of the model using a function can be straightforward. However, due to fluctuations in forage DMY caused by changing climate conditions over time, the trajectory will inevitably follow a non-linear pattern. In such cases, the fixed part of the model can be treated as factor variables, which require more degrees of freedom [38]. To address this issue, we proposed using a smooth loess function as the fixed function and selecting the polynomial order graphically (see Fig 1). By using a mathematical function instead of factors, we were able to obtain smooth trajectories over time regardless of the number of observations (see Figs. 5, 9, and 13).

### Exploring genotype by harvest interaction: reaction norms, genetic correlations, and autoregressive patterns

In forage breeding trials, the G×H interaction is a crucial aspect that needs to be understood to identify the best performing genotypes under different environmental conditions and at different stages of growth. This interaction can be studied using various statistical approaches, such as reaction norms, genetic correlations estimated by covariance functions and/or eigenfunctions across time [42].

Reaction norms are graphical representations of the relationship between genotypes and the environment. When interpretation is done by reaction norms (Figs 4, 10 and 16), it is essential to look for deviations from parallelism, such as intersections, divergences, or convergences [5]. Divergence and convergence occur when simple G×H is acting over time, meaning that the genetic variance is increasing (diverging) or decreasing (converging) [4]. The complex G×H interaction occur when the reaction norms intersect, meaning that is a lack of genetic correlation between measurements [43, 5]. In all trials analyzed, the estimated genetic correlations tended to follow an autoregressive pattern, where harvests closer to each other had higher correlations, whereas harvests far apart had lower genetic correlation (Figs. 4A, 8A and 12A). This autoregressive pattern is very common in longitudinal data in perennial species [3, 7, 19, 44, 45, 46] and have a satisfactory biological explanation, indicating that genes are expressing differently according to the environmental conditions and genotypes’ age [2, 47].

Another way to interpreting the G×H interaction in RRM is through eigenfunctions. Eigen functions are analogous to eigenvectors (principal components). Each eigenfunction is a continuous function that represents a possible evolutionary deformation of the mean yield trajectory [23]. When the eigenfunction is nearly constant, it means that the eigenfunction captured a gene pool that was equally expressed over time [23, 18, 45]. On trial T1, the first eigenfunction had a constant behavior and it is explaining the general adaptability gene pool equally expressed over time and the positive genetic correlation (Fig. 4B). The other eigenfunctions from trial T1 are explaining the lack of genetic correlation, where the gene pools had differential expression over time (Fig. 4B). For trials T5 and T7, there was negative genetic correlation over time indicating that there was not a common gene pool expressing equally through the harvests, therefore none eigenfunctions had a constant behavior (Figs. 8B and 12B). As demonstrated in this study, eigenfunctions can explain the G×H interaction as introduced by Falconer and Mackay [2], where genotype by environment interaction can be considered a pleiotropic effect of a trait evaluated across environments.

### Genotypes’ reaction norm, adaptability, and stability

A reaction norm defines a genotype-specific function that translates environmental inputs into a phenotype [5]. The differential genotypes’ response to the environment (genotype plasticity) generates the genotype by environment interaction. In this study, it was observed a higher variation for genotypes’ reaction norms for all trials interpreted (Figs 5, 9 and 13), indicating strong complex G×H interaction. Reaction norms are very informative for analyzing perennial forage over time, it allowed us to observe the behavior of each genotype, identify periods where seasonality occurred, and identify those genotypes which respond better to environmental stress. However, interpreting reaction norms for each genotype can be difficult in large breeding populations. Therefore, breeders usually use specific-genotype parameters, such as intercepts, slopes, curvatures, and variances.

These specific-genotype parameters are called sensitivity, adaptability and stability parameters in plant breeding literature and they facilitate the modeling of complex genotype by environment interactions [48-50]. Another way to select genotypes based on adaptability and stability is computing an index regarding the predicted genotypic values across all environments [18, 37, 51]. In this study, we proposed an adaptability measure the genotype’s area under the curve, which has a closer meaning to the total DMY across all harvest (*A*) and overall genotype’s performance. For stability parameter we proposed the use of the curve coefficient of variation (*CV*_*c*_). Stability can also be referred as risk in variety adoption, where the most stable genotype should have lower variance across environments, meaning that the genotype is more predictable [49, 50]. The coefficient of variation is a broad used and easy to interpret parameter in different disciplines. In time series, mainly in economics it is frequently used to infer about the risk and uncertainty in shares on the stock market exchange [52]. Therefore, we used this concept to infer about DMY stability in genotype selection, in which genotypes having lower *CV*_*c*_ will have higher stability and will also have lower variation across harvests.

## Conclusion

This study demonstrated the effective application of RRM for analyzing longitudinal data in various forage breeding trials. Our findings highlighted the importance of RRM in identifying G×H interactions and estimating adaptability and stability, using measures such as the area under the reaction norm curve and the curve coefficient of variation. By utilizing RRM in longitudinal datasets, we were able to better understand genotype seasonality through predicted reaction norms. Based on these results, we recommend the use of RRM for analyzing longitudinal traits in forage breeding trials, as it provides valuable insights and enhances our ability to interpret and evaluate genetic performance over time.

